# A numerical study about the flight of the dragonfly: 2D gliding and 3D hovering regimes

**DOI:** 10.1101/2020.03.06.980938

**Authors:** Lorenzo Benedetti, Giovanni Bianchi, Simone Cinquemani, Marco Belloli

## Abstract

The dragonfly’s ability of gliding and performing dexterous maneuvers during flight attracts the interest of scientists and engineers who aim at replicating its performances in micro air vehicles. The great efficiency of its flight is achieved thanks to the vortices generated by wing movements and thanks to the corrugations on their surfaces. The high freedom of control of each wing has been proved to be the secret behind the dragonfly capability to carry out incredible flight manoeuvers. The study presented in this paper analyzes two of the most common flight regimes of the dragonfly. Firstly, some CFD simulations of gliding are performed and drag and lift coefficients have been calculated, showing a good match with experimental data found in literature. Then, hovering has been studied using a methodology inspired to the Blade Element Momentum (BEM) theory, which is usually applied in the context of wind turbines design. The lift force calculated with this simulation corresponds to the weight of dragonfly, suggesting the correctness of this analysis.

## 1. Introduction

In the last decade bio-inspired engineering has experienced an impressive growth. The simple but clever idea behind such trend is to exploit the optimisation process carried out by Nature over thousands of years and get inspiration for realising machines that mimic the behaviour of living beings. An interesting field of application of this technique is the development of Micro Air Vehicles. MAVs are defined as “extremely small and ultra-lightweight air vehicle systems” with a maximum wingspan of 15 cm and a weight less than 20 grams [1]. Given their dimensions and their typical flight speed, which is 10 m/s at most, such “flyers” are involved in low Reynolds flight regime (i.e. from 100 to 10000), whose dynamics is ruled by a macroscopic laminal flow, affected by local turbulence. In such particular environment, force generation mechanisms are based on vortex detachment: Amalarei et al. [2] and Yao et al. [3] provided a general overview about all the vortices involved into low-Reynolds flight dynamics, whereas the majority of the articles focus on the leading edge vortex (LEV), recognised as the principal actor in this scenario ([4] - [9]).Moreover, what emerges from the definition of MAV, is that is mandatory reducing as much as possible both the payload and the space occupied by mechanical components. Under that perspective, natural flyers offer an extraordinary example to follow, with special regard to the mighty world of insects. Such reasons have boosted many researchers to study creatures belonging to this category, seeking for insights to apply in the design of MAVs. In that perspective several studies can be found about both the influence of kinematics parameters and the impact of biological characteristics on the performances of insects’ flight, mainly focusing on the effects brought in terms of forces generation and aerodynamic coefficients ([3], [10] - [16]). The contribution given by each study is specified more in detail in the following section.

On the other hand, many studies propose a CFD analysis, whose results are usually compared with those obtained by means of experimental campaigns carried out either by the same authors or by others ([3], [17] - [22]). If interested in a deeper explanation of such studies, please refer to sections 3 and 4. However, what is in common among all of them is that the proposed models are usually very complex [4]. This fact holds truer when the numerical study is extended to the 3D case [5].

Among the wide variety of insects, the scientific community recognizes the dragonfly as one of the most interesting animal to inspire the design of MAVs [6]. The reasons behind this are mainly linked to the wide range of manoeuvres that this flyer is able to perform thanks to its agility and freedom of action on the parameters controlling the flight. Such facts motivated the decision of putting the focus of this study on the dragonfly. The same choice has been made by some other researchers: Xie & Huang ([7]), Wang ([8], [9]), and [10] studied the interactions between fore- and hindwings; Kesel ([11]) and Okamoto et al. ([12]) performed experiments to investigate the effect on gliding of some relevant biological characteristics such as camber and corrugation; Tamai et al. ([13]), Mazzola ([14]), and Vargas et al. ([15]) built a CFD numerical model to simulate gliding flight.

Acquainted the importance of having a clear framework about the effect that each parameter has on the flight dynamics, this study proposes a synthetic recap of the discoveries coming from a wide range of works, together with a useful guideline for the basic understanding of low-Reynolds flight mechanisms. The truly innovative contribution of this article concerns the numerical simulations. Indeed, the present study suggests the adoption of a simplified CFD for the calculation of the aerodynamic coefficients in case of 2D gliding flight, whereas the three-dimensional hovering is studied by means of the BEM methodology. The results coming from the CFD are compared with those experimentally obtained by A. Kesel [11], pointing out a good level of agreement. For as regards 3D hovering flight it is proposed the application of BEM methodology, well established in the realm of wind turbines, instead of a much more complex 3D CFD model ([16], [5]). This new approach, proved to correctly represent the forces engendered by the flyer to keep itself aloft by developing a simple model able to catch the mechanisms of flight based on BEM. The source of interest about this paper is deemed to lie in the provision of two well-performing and relatively simple numerical tools, both 2D and 3D, with which one may test the effects on force generation brought by the parameters presented in section 2. This process should help in the proper design of experiments aimed at the realisation of MAVs.

The paper is structured as follows. A summary of kinematics and dynamics of dragonfly’s flight is presented in Section 2, where all nomenclature and reference systems are introduced. In section 3 the gliding performance are analyzed, investigating the influence on lift and drag of geometric parameters and of surface roughness in different sections of the wing and for different angles of attack, with CFD simulations. The study of hovering flight is presented in section 4, where the methodological approach is described and the results are shown. Finally, conclusions are drawn in section 5.

## 2. Flight mechanics of the dragonfly

Before entering deeper into the analysis of the flight of the dragonfly, it is considered worthwhile to guide the reader through an exhaustive introduction that allows to better understand the basis on which this work has been deployed. The flow in which the dragonfly is immersed is categorized as being in the ultra-low Reynolds number flow regime, ranging from 100 to 10000. The flapping frequency (*f*) is between 30-50 Hz.

For as regards the aspect ratio (AR), defined as the ratio of the wingspan to the wing mean chord AR = b/c, the one of the dragonfly typically ranges from 8.4 to 11.63 [17]. The higher the aspect ratio, the higher the lift-to-drag ratio. The front of fore and hind wings has longitudinal and transversal veins, whose function is to stiffen the thin membranous wing so that the insect can withstand the inertia forces acting on it [18]. In contrast with the common knowledge about well-designed airfoils, there is proof that under the flow conditions in which dragonflies are involved, such structure enhances the performances of the wing [12] [11].

Wing muscles insert directly at the wing bases, which are hinged so that a small movement of the wing base downward, lifts the wing itself upward, very much like rowing through the air [14] [9]. Dragonflies have fore and hind wings similar in shape and size, each of which operates independently giving a degree of fine control and mobility in terms of the abruptness with which they can change direction and speed.

The wing tip traces a figure-8 pattern with respect to the wing root [19] [20]. In one flap cycle, the wing revolves around the hinge back and forth at roughly constant angle of attack, while rotating along the spanwise axis during stroke reversal at the beginning and end of a stroke. When rotating from down to upstroke, the morphological underside of the wing rotates to face upwards and this is called supination, whereas from up- to downstroke the morphological underside of the wing rotates to face downwards again, and this is called pronation [18].

**Figure 1:**
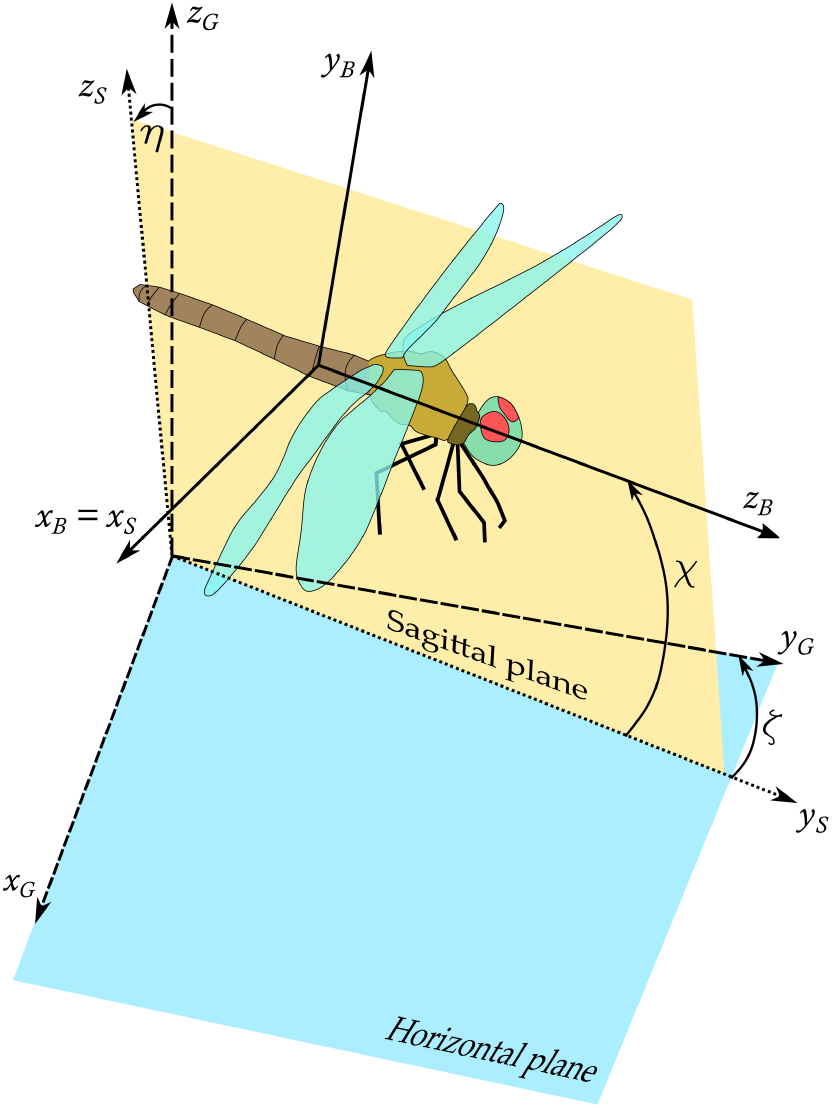
Geometric model of the insect with affixed body frame (*x, y, z*)^*B*^ in global reference frame (*x, y, z*)^*G*^: *ζ, χ*, and *η*. denote the yaw, pitch, and roll angles respectively

**Figure 2:**
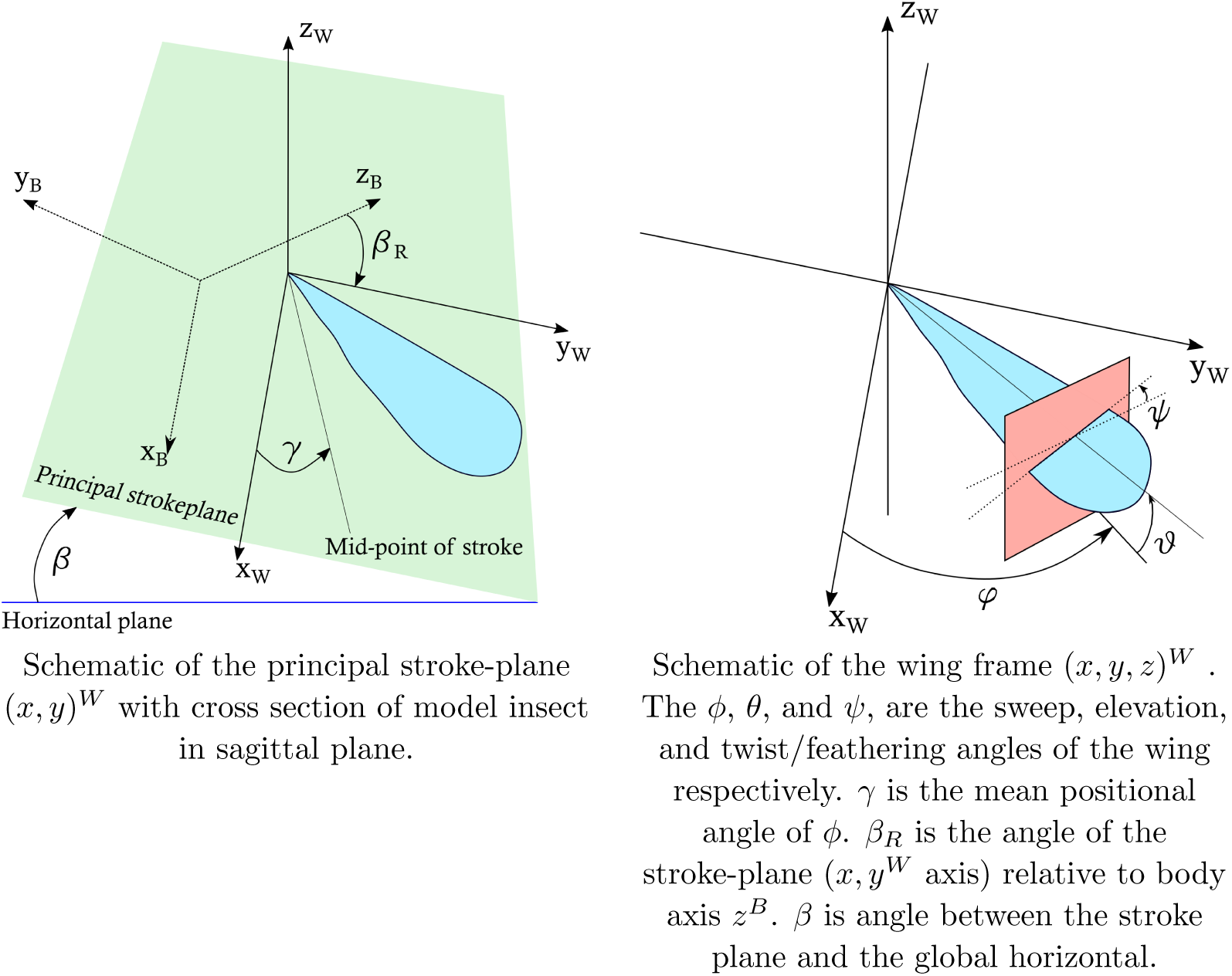
Angles of the wings

**Figure 3:**
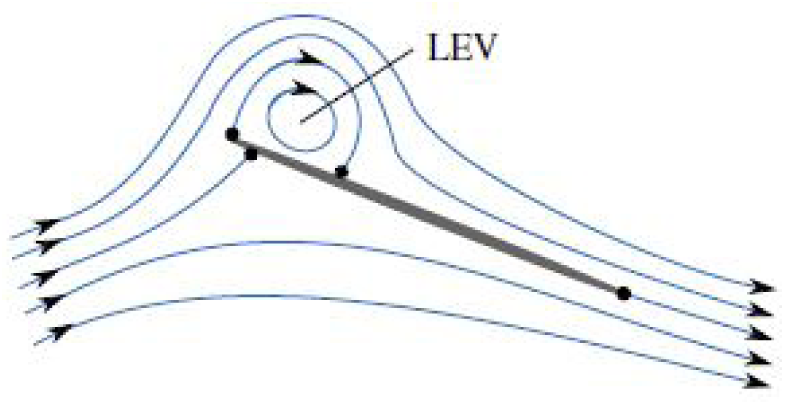
Schematic showing the simplest valid LEV structure for a cylindrical vortex-cross-section view. The LEV is stable at high angles of attack with flow reattachment on the upper surface. Black dots represent stagnation points

**Figure 4:**
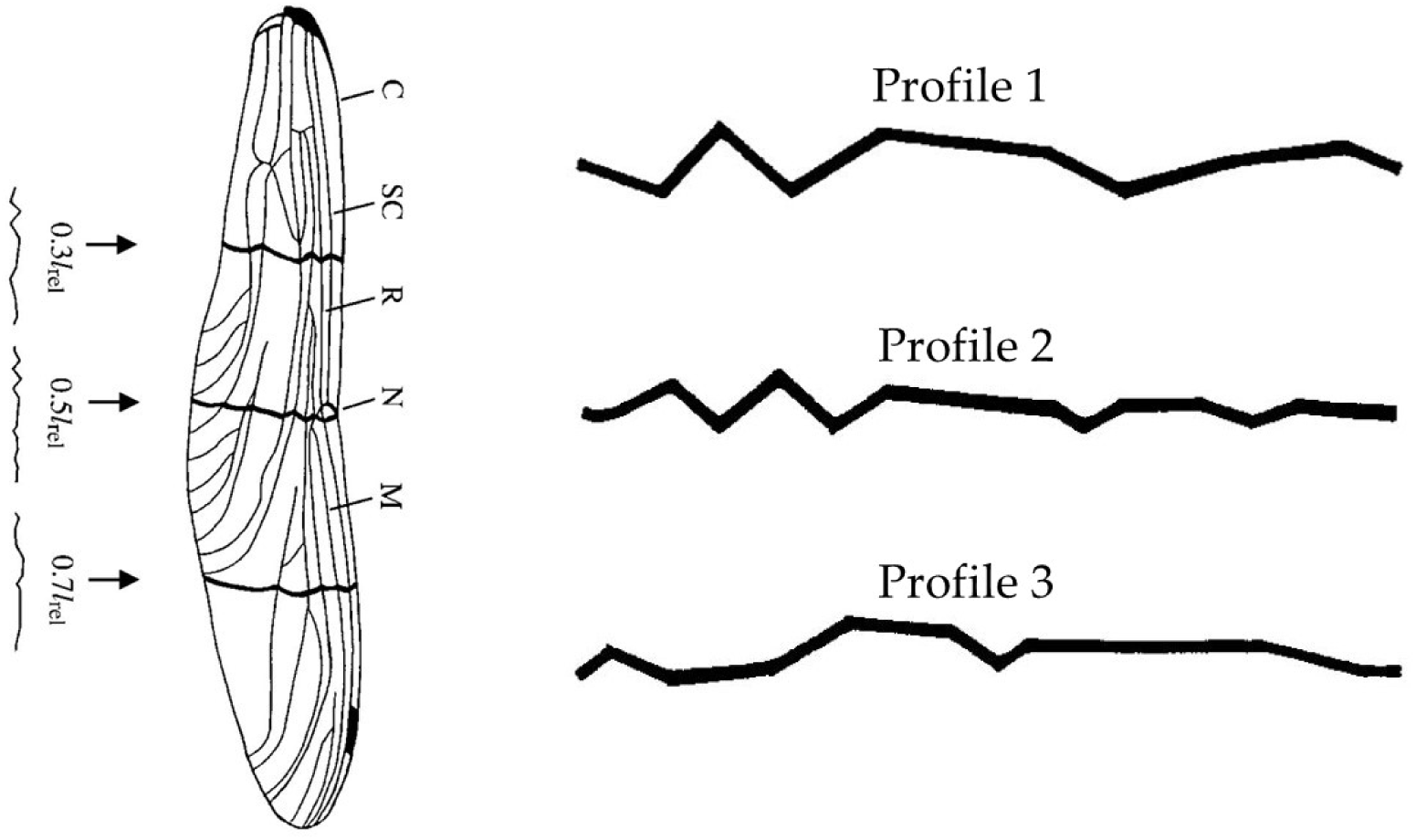
Set of profiles used for the simulations. On the left: representation of the whole wing, with indication of sectioning locations. On the right: the three profiles, from up to down, corresponding to sections 1, 2, and 3, respectively. Source:[11]

**Figure 5:**
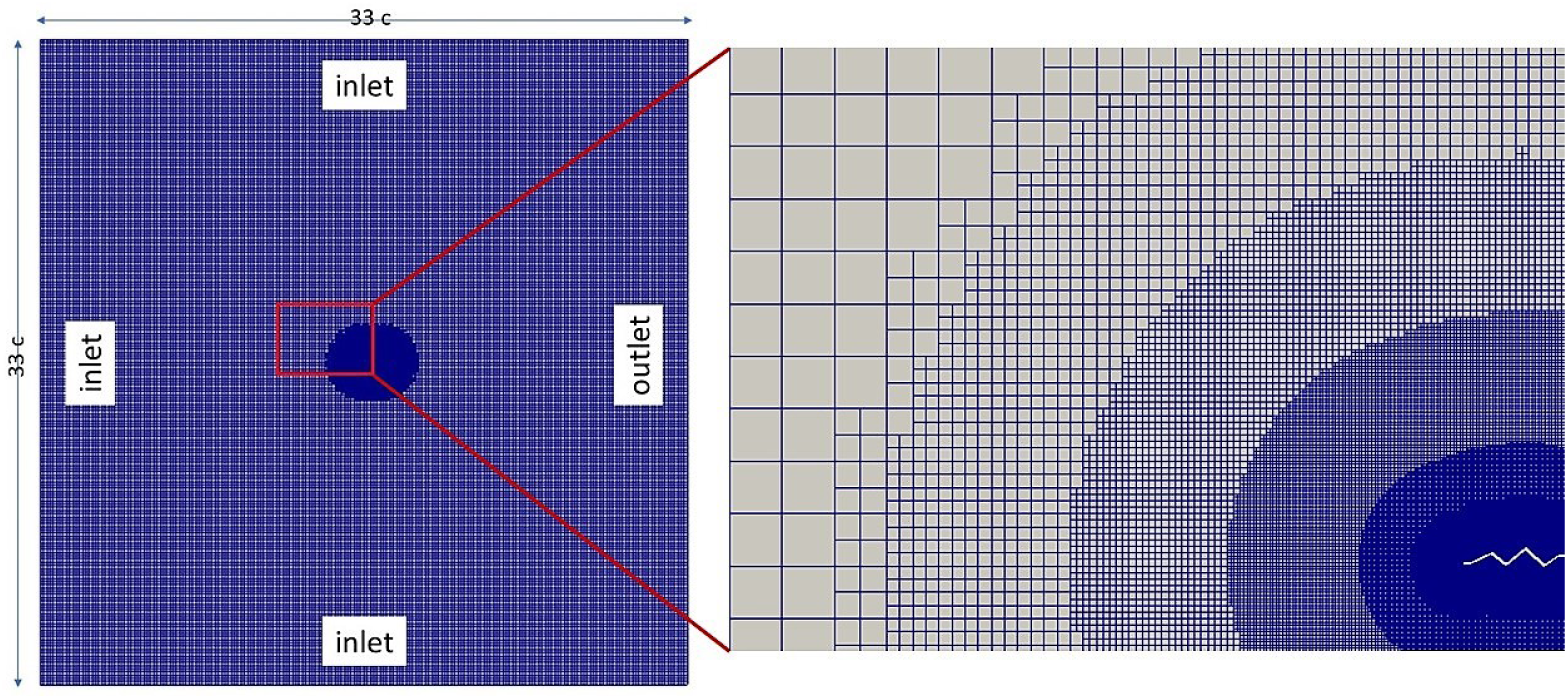
On the left: view of the complete mesh. The tags in correspondence of the four sides indicate the type of boundary condition applied to each of them. The dimensions of the grid are intended as multiples of the chord length *c*. The central red box identifies the zooming in proximity to the surface, represented in the image on the right. On the right: Detail of the portion of the mesh enclosed in the red box. 6 refinement levels should be identified, from the background mesh to the vicinity of the profile

**Figure 6:**
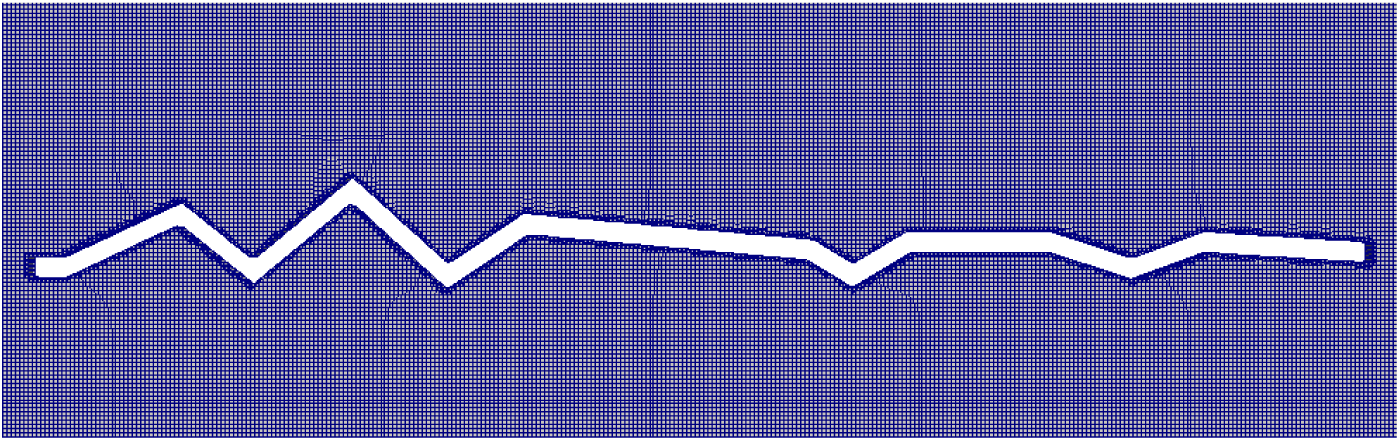
Framing realised zooming on the profile. The last refinement level may be noticed looking at the cells the closest to the surface of the wing. No boundary layer is present

**Figure 7:**
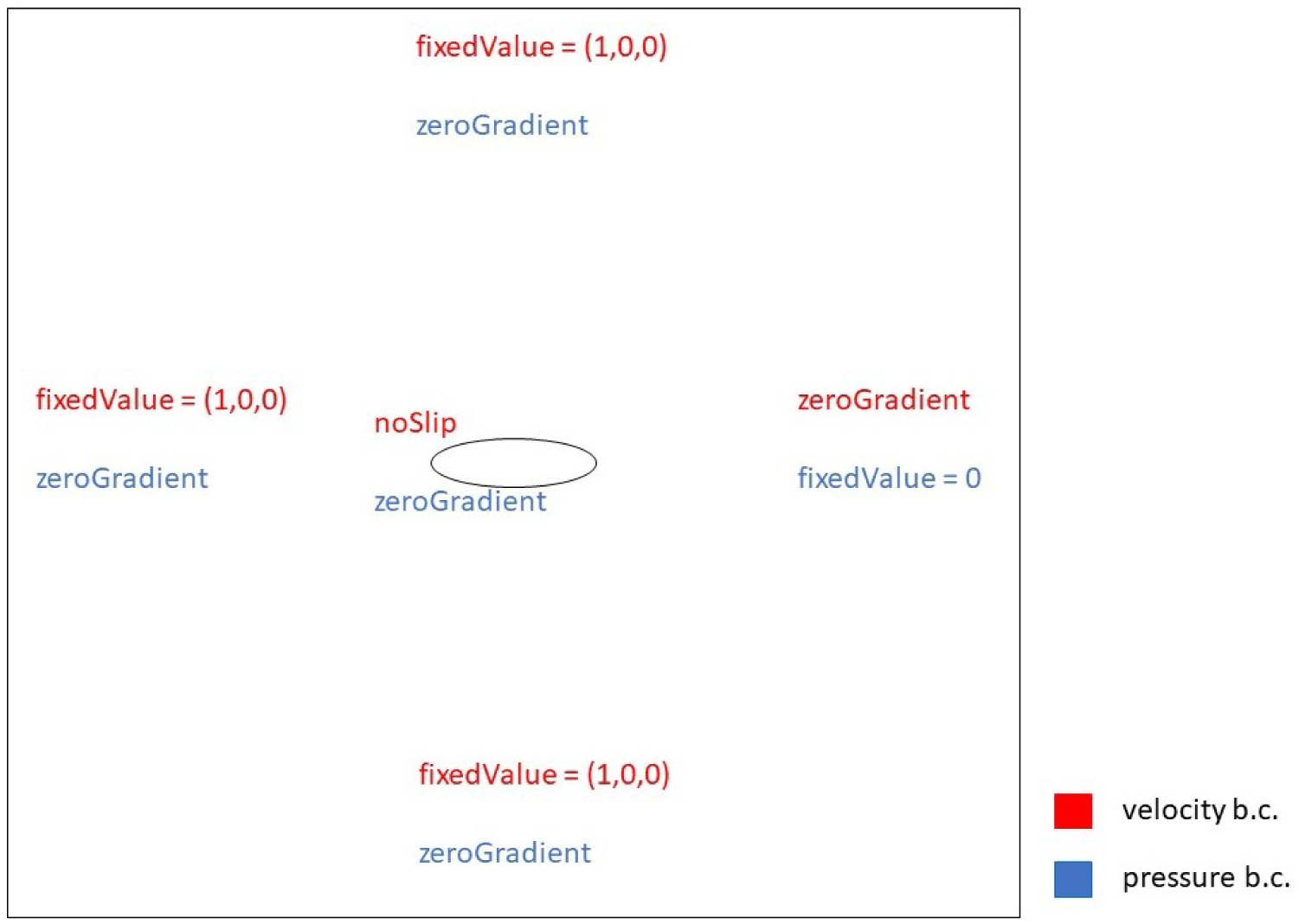
Representation of the boundary conditions assigned to velocity and pressure fields, in correspondance of each patch. On the bottom right corner the legenda is provided. This framework is referred to AoA = 0° cases.

**Figure 8:**
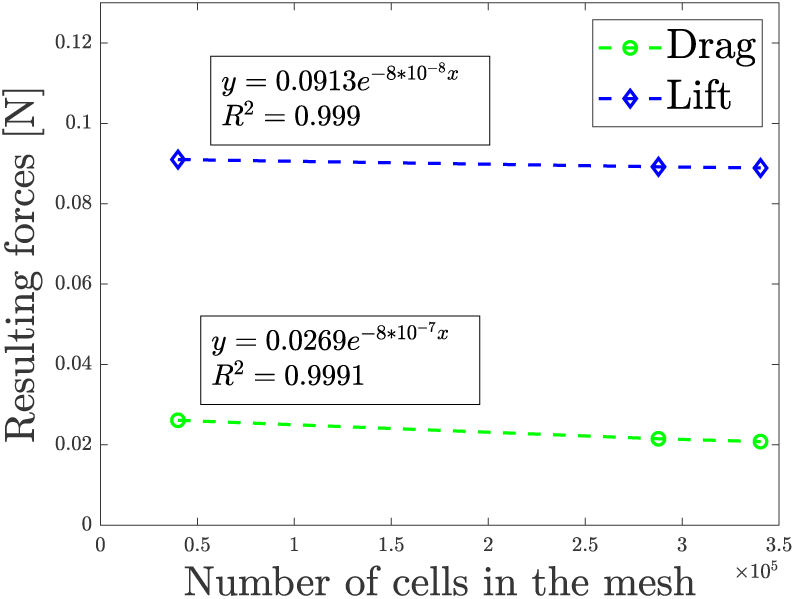
Trends of lift and drag forces, together with the exponential fitting and the related *R*^2^ factor.

The flap cycle can be modelled as a sinusoidal motion [18]:

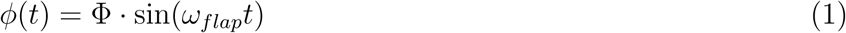

Henceforth, Φ will be called sweep angle, while *ϕ* represents the stroke amplitude. Both parameters are extensively dealt with when the kinematic framework is presented. Intuitively, a wing sweeps an angle equal to 2Φ per cycle. To define an average Reynolds value, constant over the whole flapping cycle, the motion must be simplified considering it as if it were characterized by a constant angular velocity. Under that hypothesis, a reference wingtip velocity can be determined: *U*_*ref*_ = *R ·* 2Φ *· f*. As a consequence, the Reynolds number may be defined as:

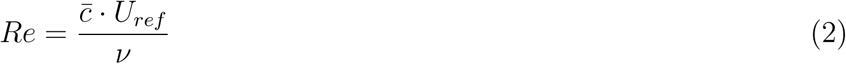

Here,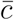 is the mean wing chord, and *v* is the kinematic viscosity of air (1.5 *·* 10^−5^ *m*^2^*s*^−1^). The small size of the flyers and the relatively low cruise speed they sustain, cause the Reynolds number to hardly go beyond 10^3^. Such condition of motion belongs to the so-called low Reynolds flight regime and, although the dragonfly is one of the biggest insects and surely among the fastest ones, its flight still develops into a purely laminar flow domain.

Exploiting the just defined reference velocity, the advance ratio (*J*) can be introduced:

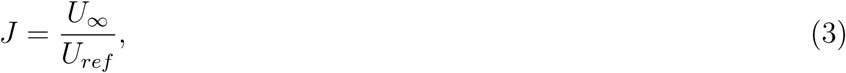

where *U*_*∞*_ and *U*_*ref*_ represent the velocity of the insect and the wingtip velocity, respectively. To better describe the flight, let us define two different cartesian reference frames: a global one (*x, y, z*)^*G*^, which is fixed at ground, and a local one (*x, y, z*)^*B*^, rigidly attached to the animal’s body. The orientation of the model example with respect to the fixed frame is specified by the angles of yaw (*ζ*), pitch (*χ*), and roll (*η*) [3] (Figure 1).

The principal stroke plane (*x, y*)^*W*^ is shown, and the wing frame (*x, y, z*)^*W*^ is defined as for the body and the ground. The kinematic link between the reference system of the body and the one of the wing is represented by the alignment between *x*^*B*^ and *x*^*W*^ [3].

The first angle to be defined is the stroke-plane angle *(β*). The stroke-plane angle is an approximate measure of the relative angle of the stroke plane (*x, y*)^*W*^ to the global horizon. Then, three more angles are crucial to describe the motion of the flyer: the main flapping motion or sweep angle Φ, the slight deviation from the stroke plane or elevation angle *θ*, and the active wing rotation or twist angle Ψ [3].

Aerodynamic forces should be considered as to be applied in the so-called center of pressures. However, its location moves significantly with a change in angle of attack and is thus impractical for analysis: the most adopted solution is to exploit the so-called aerodynamic center. It is a common choice to put the AC in correspondance of the quarter-chord point, which is located approximately 25% of the chord from the leading edge of the airfoil.

An important key for the comprehension of insect flight is the understanding of the vortex shedding due to a rapidly oscillating wing. The presence of a leading-edge vortex (LEV) generated on top of the flapping wing, increases the lift force to values much higher than those predicted by conventional wing theory [17]. The capture and recirculation of the LEV has been extensively studied as an unsteady mechanism primarily responsible for generating lift over insect wings. It consists of a vortex with appreciable spanwise flow maintained at the leading edge of the wing during the upstroke and downstroke. At pronation and supination, the vortex is shed, after which a new vortex is quickly formed [21]. Several studies have shown that the wing benefits from the attachment of the LEV because of the low-pressure core of the LEV acting on the wing. The origin of the leading-edge vortex is the roll-up of shear layers, present in highly viscous flows, which is the case at Reynolds numbers as low as the ones characterizing insects’ flight. Such vortex is stable because its location remains near the leading edge and it does not grow with time [22]. As shown in figure 3, this allows the flow over the upper surface of the wing to separate at the leading edge but then reattach before the trailing edge.

The wing translation creates a pair of leading and trailing-edge counter-rotating vortices, while the wing rotation then combines them into a dipole. Thanks to their mutual induced flow, they form a comoving pair. If the two movements are properly combined in terms of phase, the dipole moves downward carrying momentum with it to generate a lift on the wing. Eventually, the self-induced flow sweeps away the vortices from the wing, so they do not interfere with the vortices created by the subsequent cycle. Such mechanism reveals that the figure-8 motion manages to solve two problems simultaneously: to create the vortices dipole and to get rid of it [23].

Hereafter the influence of the main biological and kinematic parameters is summarized:

- **Leading-edge sharpness**: being the leading edge the first part of the wing to get in touch with the airflow, the sharper it is the easier it is to “cut” the fluid in which it is immersed. Leading-edge sharpness promotes flow separation, facilitating the formation of the LEV especially at low angles of attack, in correspondence of which it is more difficult to create vortices[24] [12].
- **Camber**: its effect is noticeable especially during gliding, in particular a negative curvature (i.e. downward convex profile) is detrimental both in terms of *C*_*L*_ and *C*_*D*_, while a positive one is desirable, since the lift coefficient increases far more than the drag one [11]. Also, the maximum lift-to-drag ratio (*L* = *D*)_*max*_ attained by an upward convex profile is higher than the one pointed out by the competitors, leading to an increased efficiency. For gliding performances, when the location of the maximum camber is fixed, increasing the value of the latter will result in greater lift, and this effect is more significant at large AoAs, at the same time the drag is slightly influenced [12].
- **Corrugation**: comparing a corrugated airfoil with a streamlined profile and a flat plate immersed in a low Reynolds number flow, the outstanding performances of the former become evident, being higher the angle of attack at which stall begins. The unsteady vortical structures shedding from the protruding corners were found to be trapped in the valleys of the corrugated cross section, which dynamically interact with the high-speed flow streams outside the valleys. Thanks to the interaction between the unsteady vortex structures and outside high-speed flow streams, high-speed fluid is pumped to near-wall regions, which provided sufficient energy for the boundary layer to overcome the adverse pressure gradient to suppress flow separation and airfoil stall [12] [11] [25] [13].
- **Flapping frequency(***ω*): this parameter is confined, in the specific case of the dragonfly, between 30 and 50 Hz, and it has been demonstrated that the higher the frequency, the higher the lift coefficient. In accordance to that, it has been registered a significant increase of the flapping frequency in the initial phase of acceleration, followed by a stabilization around a steady-state condition [3].
- **Mean positional angle (***γ*): the main techniques exploited to reach a certain speed, such as paddling and pitching, are responsible for the generation of an important pitch-up moment that needs to be at least diminished. An important and natural counter action used by insects is obtained by merely biasing the mid-point of the wing strokes rearward (dorsally) via the mean positional angle. Biasing the sweep of the wings backwards effectively moves the center of mean wing force rearward relative to CoM, thus inducing an increase in pitch-down moment on the model insect to counter the pitching-up tendency. Similarly to the flapping frequency, also biasing of the sweep angle is greater in the initial phase of the flight, when the pitch-up moment is more severe, successively it is stabilized to a lower value. In case of backward flight things work in an opposite way [3].
- **Elevation angle (***θ*): This degree of freedom is one of the most peculiar ones, since it gives rise to the famous “8-shaped” wingbeat [26]. This parameter has a twofold impact on the flight dynamics: it is useful to counterbalance the pitch-up moment acting on the flier, and it is also important to level the force distribution during the wingbeat [27] [3]. It is able to provide an amount of pitch reduction comparable to the mechanism of regulating the mean positional angle and it may be exploited alongside with the latter to stabilize the flight. The reason behind these outcomes is that the elevation considerably increases the effective angle of attack just after the stroke reversal, enhancing the LEV [27].
- **Stroke plane angle (***β*): this kinematic parameter is linked mainly to the amount of thrust generated, in particular, for any given airspeed *V*_1_, the forward tilt of the stroke plane increases the thrust and decreases the lift; for a given *β*, the thrust decreases with increasing airspeed, eventually becoming a drag at sufficiently high airspeeds. The lift force, on the other hand, generally increases with the airspeed. A larger forward tilt of the stroke plane is thus required to sustain a higher forward velocity [3].
- **Angle of attack (***α*): it plays a substantial role in governing the LEV [28], and an asymmetry of the AoA between the up and the downstroke can effectively produce mean thrust force to propel the flier in a range of airspeeds even when the stroke-plane angle is horizontal, this motion is called “paddling” and for any given airspeed *V*_*∞*_ it increases the thrust and decreases the lift. For a given asymmetry of AoA Δ*α*, the thrust decreases with increasing airspeed, eventually becoming a drag at sufficiently high airspeeds; that paddling produces a drop in the corresponding mean pitch-up moment on the insect. This is desirable as it could offset pitch-up torque associated with the use of stroke-plane tilting to generate thrust, and hence reduce the burden of moment/torque regulation during flight. This synergy between the two techniques encourages fliers to combine them to attain higher speeds [3].

## 3. Gliding: numerical analysis

Gliding is a particular flight condition in which the insect does not beat the wings, still being capable to move forward. Such maneuver is not so common among insects: indeed only some species, including the dragonfly, have sufficiently large wings to sustain it. Notwithstanding, the study of gliding is of great interest since the majority of MAVs are provided with wings that do not beat. This analysis starts from the approach followed by A. Kesel [11], where three profiles obtained sectioning a forewing of an *Aeshna Cyanea* are assessed in terms of lift and drag coefficients. The three sections of the wing considered for this analysis are shown in figure 4, the exact geometry of the corrugations is taken from [11] and it has been replicated in a CAD model.

The assessment of the results is made comparing the calculated lift and drag coefficients with those obtained experimentally in [11], keeping the same Reynolds number of 10000.

Each section is analyzed independently with 2D steady-state simulations using OpenFOAM v5. The mesh is obtained with 6 refinement levels, as shown in figure 5, keeping unchanged the mesh of the domain while changing the direction of the relative velocity. This is made possible by declaring three edges as “Inlet” as shown in figure 7. Such decision implied the necessity of having a grid large enough to attain a perfectly undisturbed flow at the boundaries, to avoid any possible unrealistic rebound of the flow.

Once is chosen the type of the mesh, a sensitivity analysis has been performed to select the correct number of cells. Exploiting a trial&error process, comparing two successive sets of mesh a difference of less than 5% was tolerated with respect to drag force, while a more precise evaluation of the lift force has been required, with the threshold level set at 1%.

Among the wide range of solvers available in OpenFOAM, SimpleFoam was chosen for running the cases treated in this section. SimpleFoam is a steady-state solver for incompressible, turbulent flow. The focal features of this solver, with the aim of understanding why it has been deemed to be the most appropriate for this work are listed below:

- **Turbulent**: Even if this might seem in contrast with the low Reynolds realm in which dragonflies live, the particular geometry of the profiles under analysis, featured by many severe pleats acting as turbulators, causes the born of many vortices whose structure is better appreciated by a turbulent flow solver;
- **Incompressible**: dealing with air, this condition is always respected with flow velocities under 100*m/s*;
- **Steady-state**: the nature of the dragonfly’s flight is characterized by some important unsteady effects [51]; nevertheless, they appear mainly during flapping and they can be considered negligible during gliding.

As far as turbulence is concerned, Navier-Stokes equations have been solved focusing on the average flow, according to the Reynolds-averaged-Navier-Stokes (RANS) method. The adopted turbulence model used is k-*ω*SST, with the following values of its parameters.

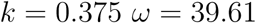

### 3.1. Analysis of fluxes

To consider different flight conditions, for each profile shown in figure 4 many simulations have been carried out for different angles of attack ranging from −5° to 10°. Beyond such value the second profile taken into account by Kesel suffers from incipient stall [13]. For each profile only the results for angles of attack −2°, 0° and 2° are commented for the sake of brevity. Further increasing the AoA the flow topology evolves according to the same trend. The profile in the center of the wing is the most interesting to analyze deep because of its high consideration in literature [14], [13], [15]. For the other analyzed profiles the results are presented just in form of pictures; making the same considerations made about profile 2, the reader can easily understand how forces are generated in different conditions.

Looking at the shape of profile 2, shown in figure 4, it can be noticed that the leading edge is almost horizontal, while the trailing edge slightly faces downward. In figure 9 the results of the simulations of this profile with AoA of 0° are presented.

**Figure 9:**
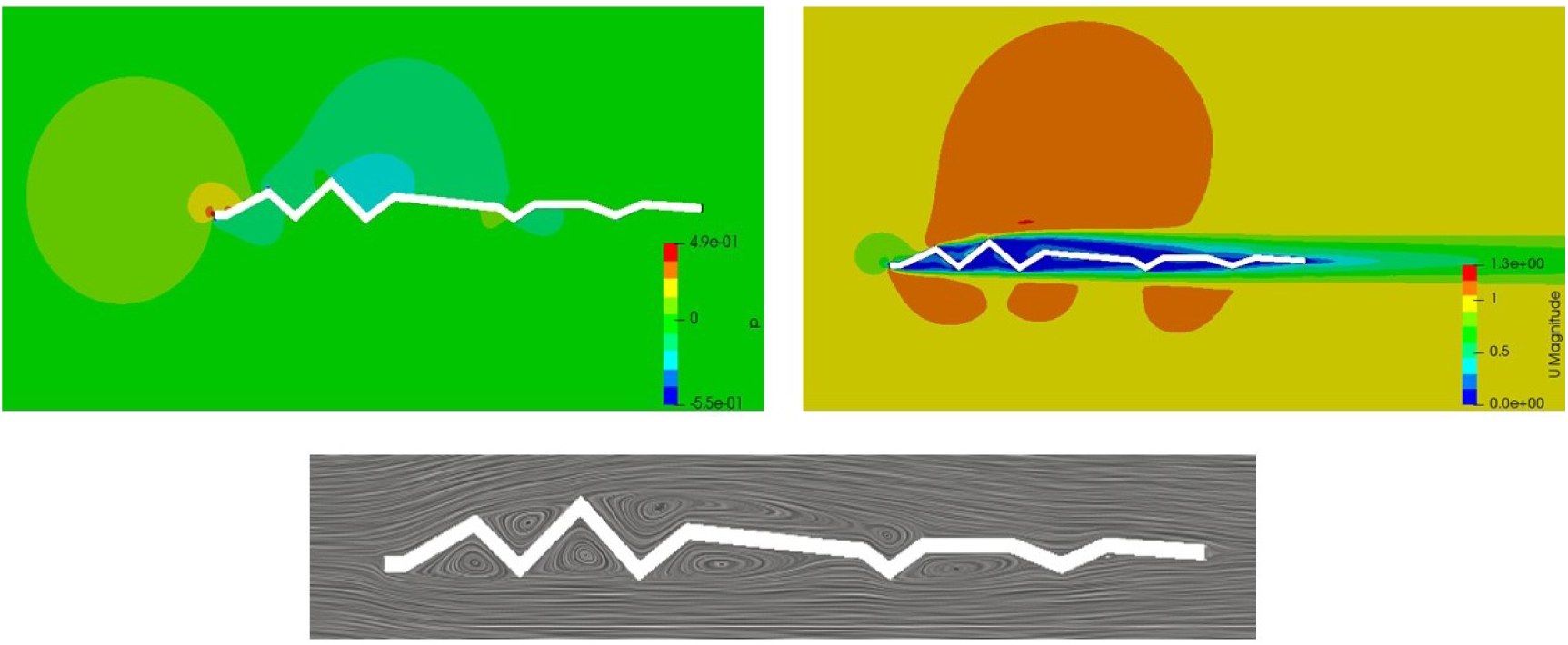
Up, left: pressure field. Up, right: velocity field. Bottom: streamlines of the vortices. Results referred to profile 2, in correspondence of an AoA = 0°

It is possible to notice a high-pressure region exactly in front of the leading edge, which does not equally expand towards the two sides of the wing. Conversely, it is more oriented to the upper part, due to the more favorable inclination pointed out by the first up-edge compared to its down-parent. The exact same shape is reproduced by the velocity perturbation. Regarding the bottom side, the shape of the first pleat just after the leading edge, less smooth than the one on the upper side, causes the first flow separation, soon replicated by the following edges, in correspondence of which a reattachment with subsequent detachment takes place. The result is a series of three perturbed zone not big enough to merge. With such velocity field, it should be now easy to imagine the configuration of the pressures: a big low-pressure region above the wing should be expected, as confirmed by the image on the left, whereas three smaller depressions are found below the profile, the most important one nested in the first pleat just after the leading edge. The dynamic translation of such pressure distribution is a slightly positive lift force generation, favored by the big underpressure above the middle of the profile despite a bit of overpressure witnessed near the leading-edge.

The pressure distribution and the air flow with an AoA of −2° are shown in figure 10. Looking at the pressure distribution, the effect of the airflow direction is immediately visible: a perturbed region, evidences an overpressure in correspondence of the leading edge, upper side. In parallel, a decrease in velocity may be noticed. Regarding flow separation, two big zones, one above and one below the profile, may be identified. The first one occurs when the fluid interacts with the first edge after the leading one, whose neutral direction cannot promote any detachment in case of negative angle of attack. Looking to the bottom side, the relative angle between the leading edge and the air flow is sufficient to cause flow separation, bringing an intense perturbation in the vicinity of the leading edge, diffused then to all the area under the profile. This situation, described looking at the velocity field, is clearly reflected also by the pressure distribution: two significant areas are highlighted, one nested inside the first pleat of the bottom side, characterized by a severe depression, and a less intense one in correspondence of the second valley, upper side. Around such regions, two other perturbed area can be visualized, the one underneath the profile is of far bigger extent. In light of these considerations, a negative lift force is pointed out.

**Figure 10:**
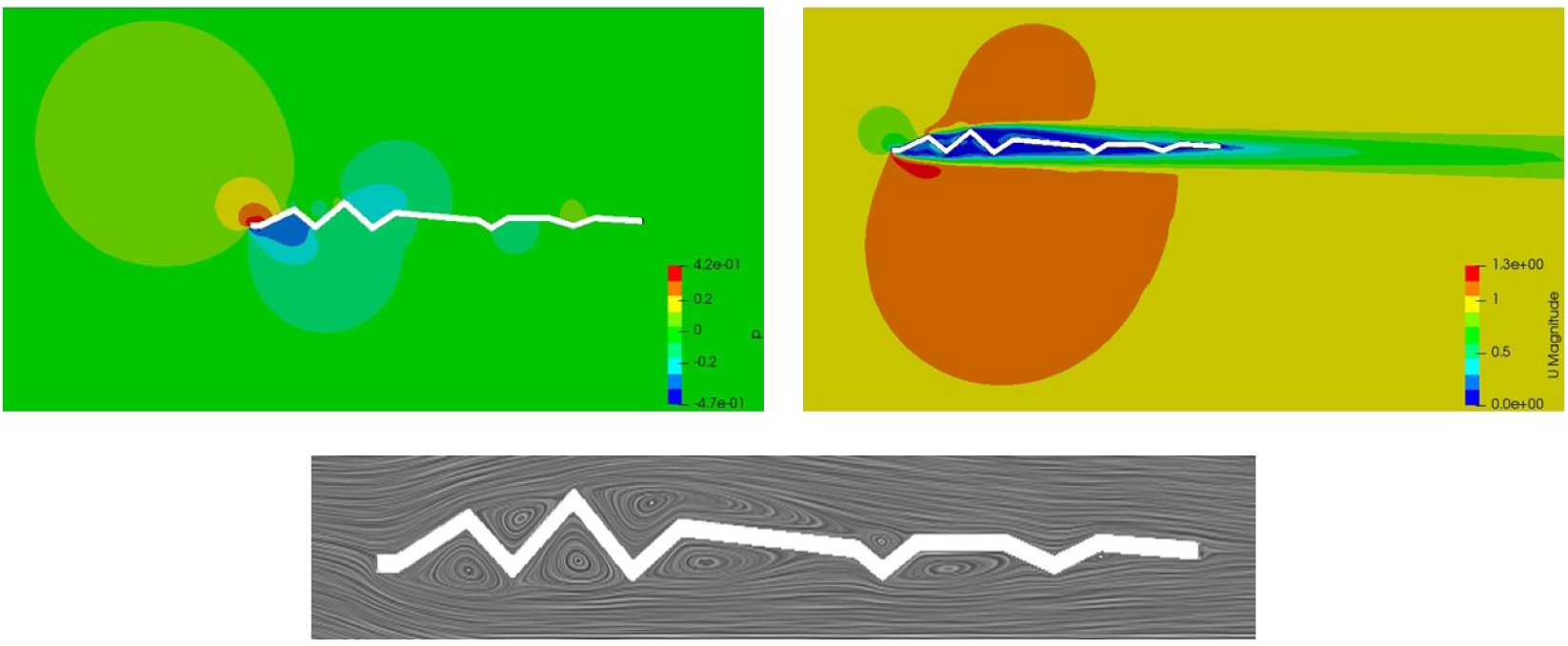
Up, left: pressure field. Up, right: velocity field. Bottom: streamlines of the vortices. Results referred to profile 2, in correspondence of an AoA = −2°

The results regarding this profile with an angle of attack of 2° are shown in figure 11.

**Figure 11:**
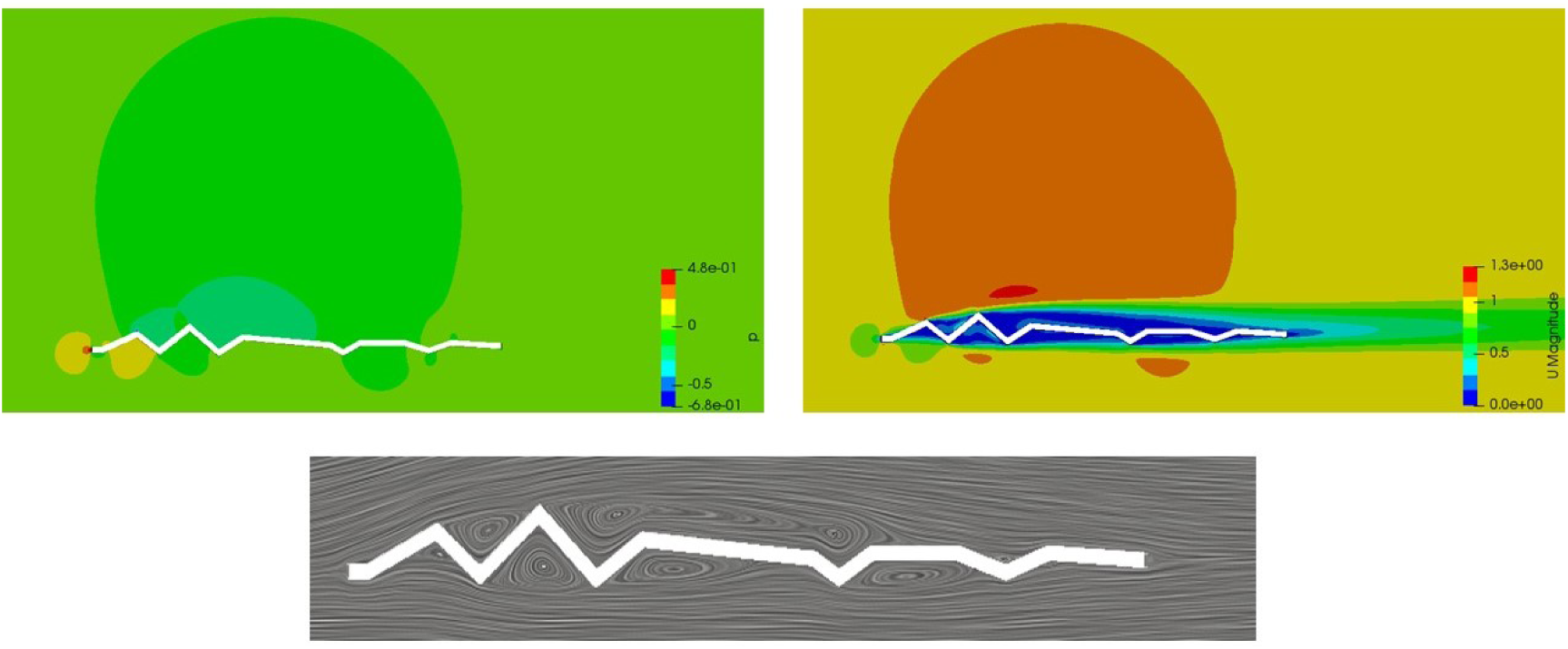
Up, left: pressure field. Up, right: velocity field. Bottom: streamlines of the vortices. Results referred to profile 2, in correspondence of an AoA = 2°

Increasing the angle of attack from 0° to 2° the flow topology evolves coherently. Indeed, the high-speed region above the profile extends in dimension, still keeping the maximum value to 1.3 m/s. Also the position of the peak holds still, in correspondence of the second edge. Looking to the lower side, the opposite behavior is pointed out, with high-velocity zones that almost vanish. The null inclination of the leading edge, together with the unfavourable shape of the first pleat below the wing make the circular area to split into two parts, one confined to the front of the profile, the other one completely filling the first valley underneath. This fact clearly impacts on the vorticity field: the recirculation zone inside the first pleat almost disappears, leaving space to a slowdown of the air flow until reaching stagnation in the very proximity to the wing. The pressure field adapts accordingly: two overpressurized areas form, one ahead the profile, one nested in the subsequent valley; furthermore, a huge depression region is born above the wing, strongly contributing to an important lift generation.

A negative lift force for an angle of attack lower than 0° and a positive lift force for positive angles of attack is observed for every analyzed profile. Due to the different geometry of the corrugations of the profiles, the generated vortices have different positions and strength; however, the mechanism underlying their role in lift generation is exactly the same and it can be observed in figures from 12 to 17.

**Figure 12:**
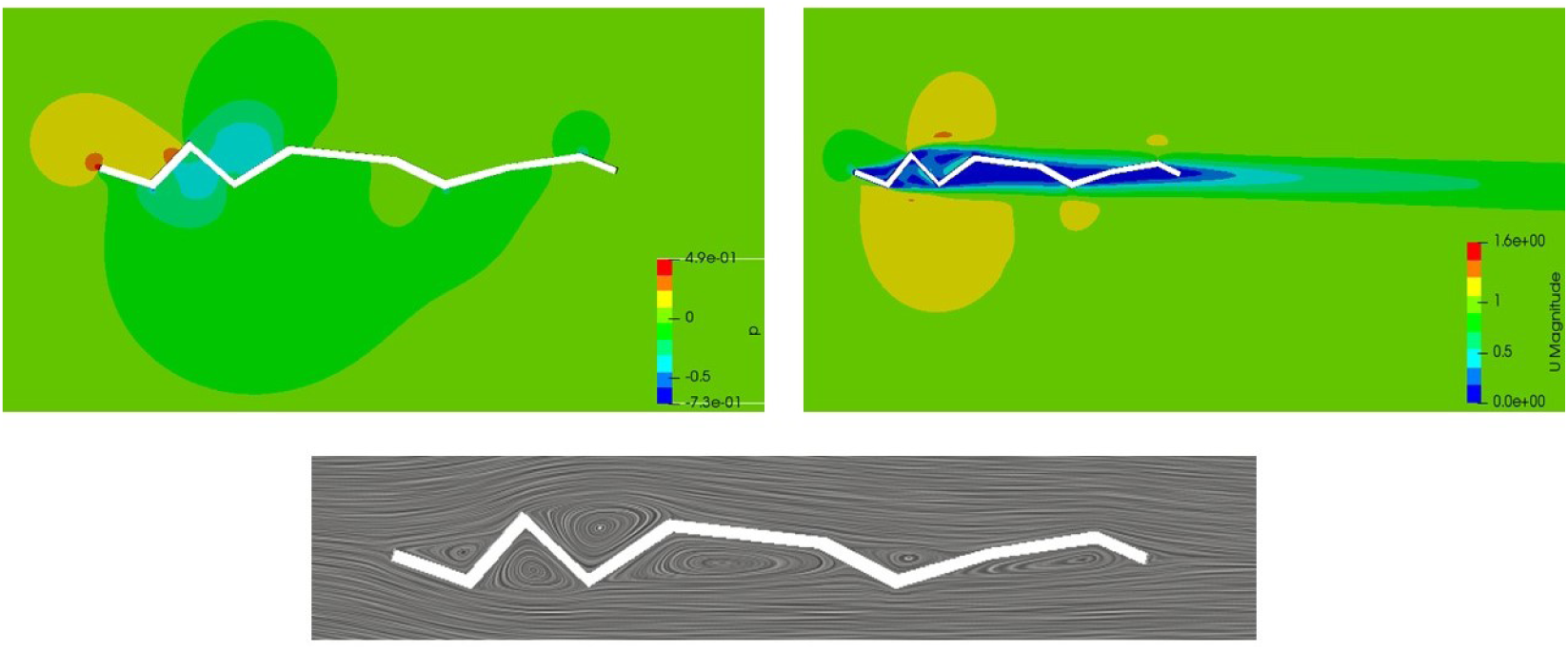
Up, left: pressure field. Up, right: velocity field. Bottom: streamlines of the vortices. Results referred to profile 1, in correspondence of an AoA = −2°

**Figure 13:**
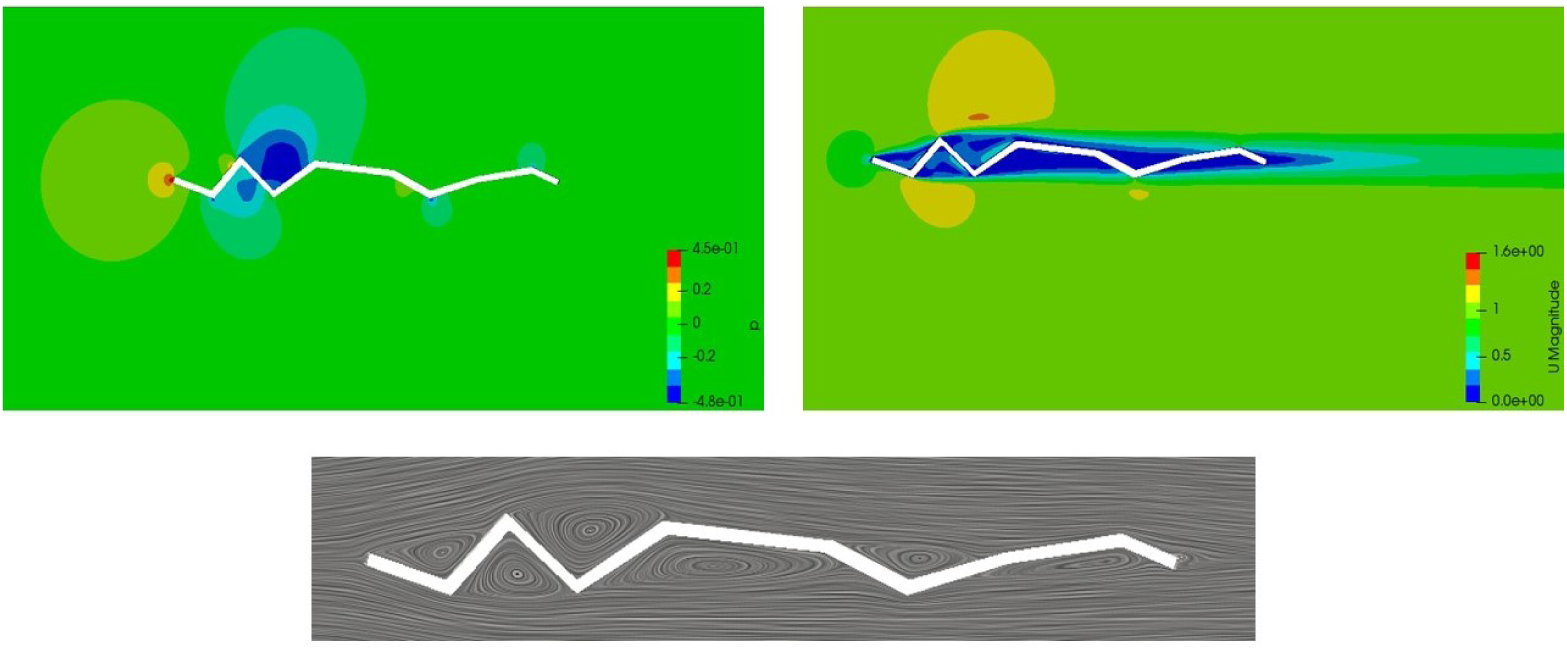
Up, left: pressure field. Up, right: velocity field. Bottom: streamlines of the vortices. Results referred to profile 1, in correspondence of an AoA = 0°

**Figure 14:**
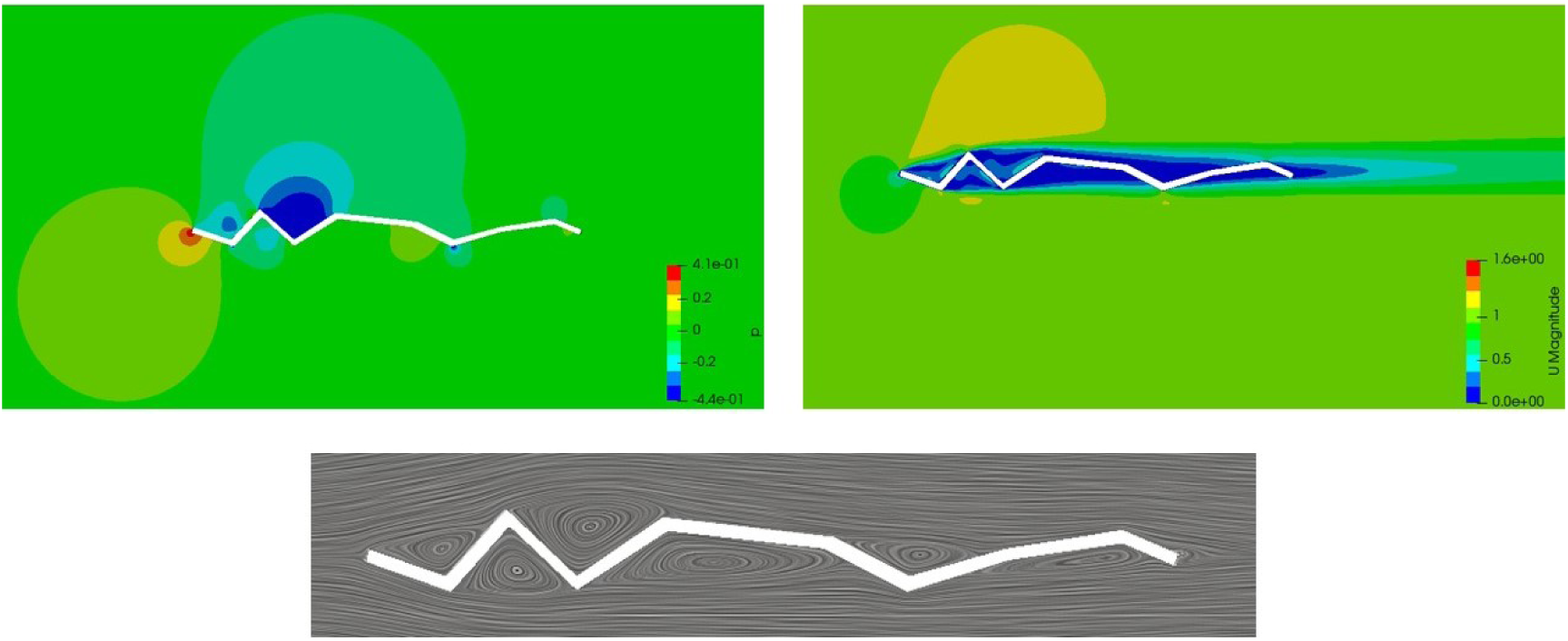
Up, left: pressure field. Up, right: velocity field. Bottom: streamlines of the vortices. Results referred to profile 1, in correspondence of an AoA = 2°

**Figure 15:**
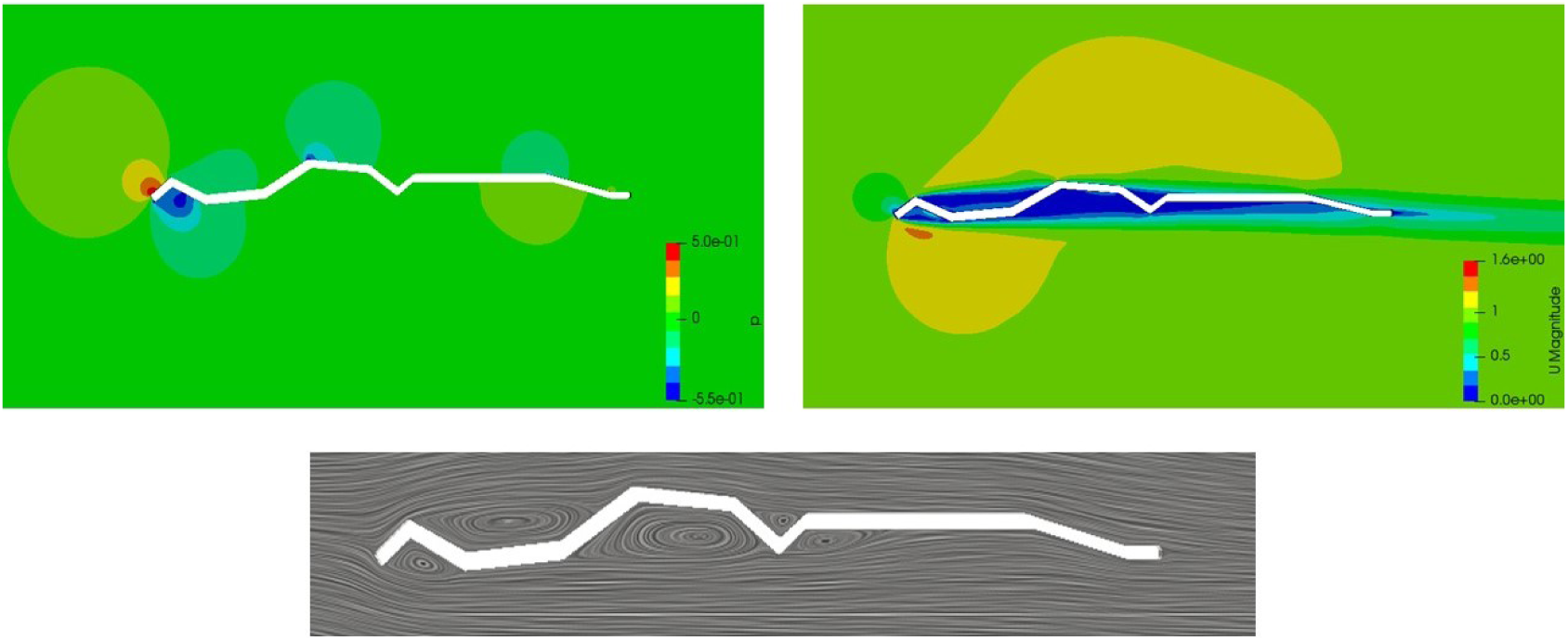
Up, left: pressure field. Up, right: velocity field. Bottom: streamlines of the vortices. Results referred to profile 3, in correspondence of an AoA = −2°

**Figure 16:**
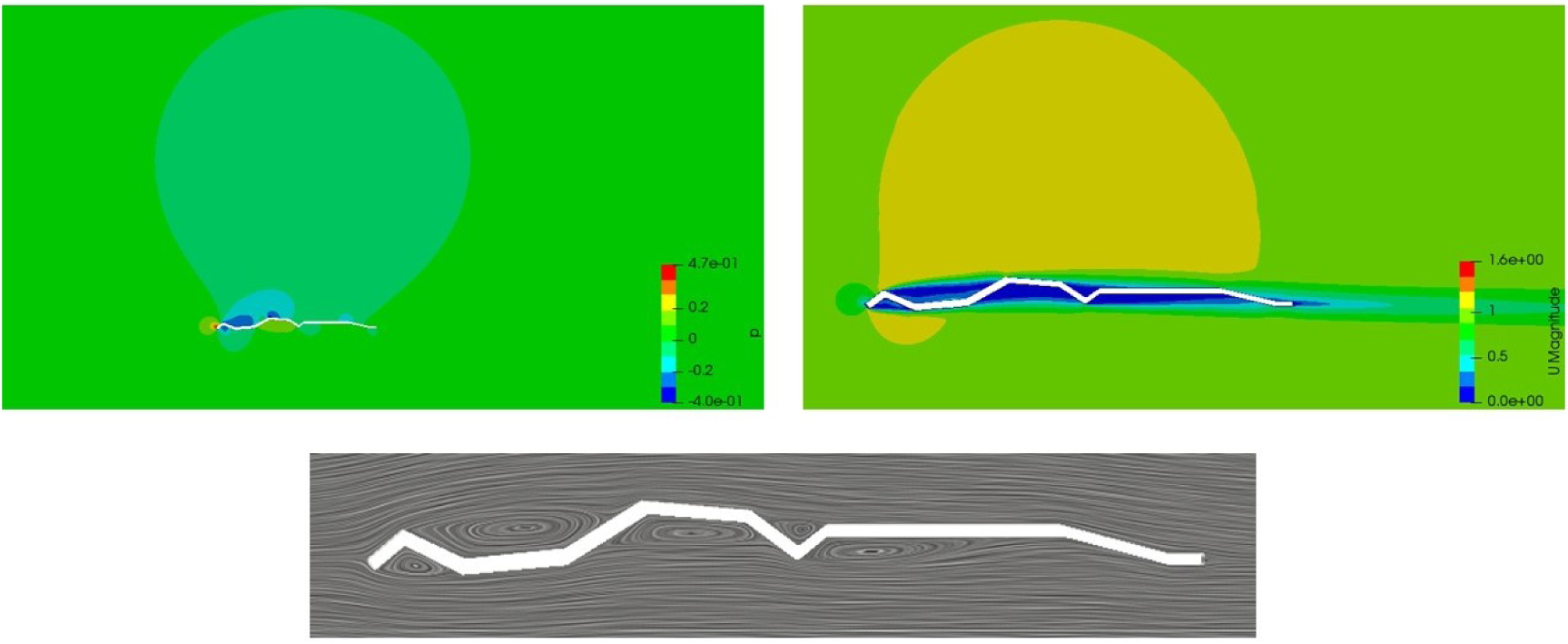
Up, left: pressure field. Up, right: velocity field. Bottom: streamlines of the vortices. Results referred to profile 3, in correspondence of an AoA = 0°

**Figure 17:**
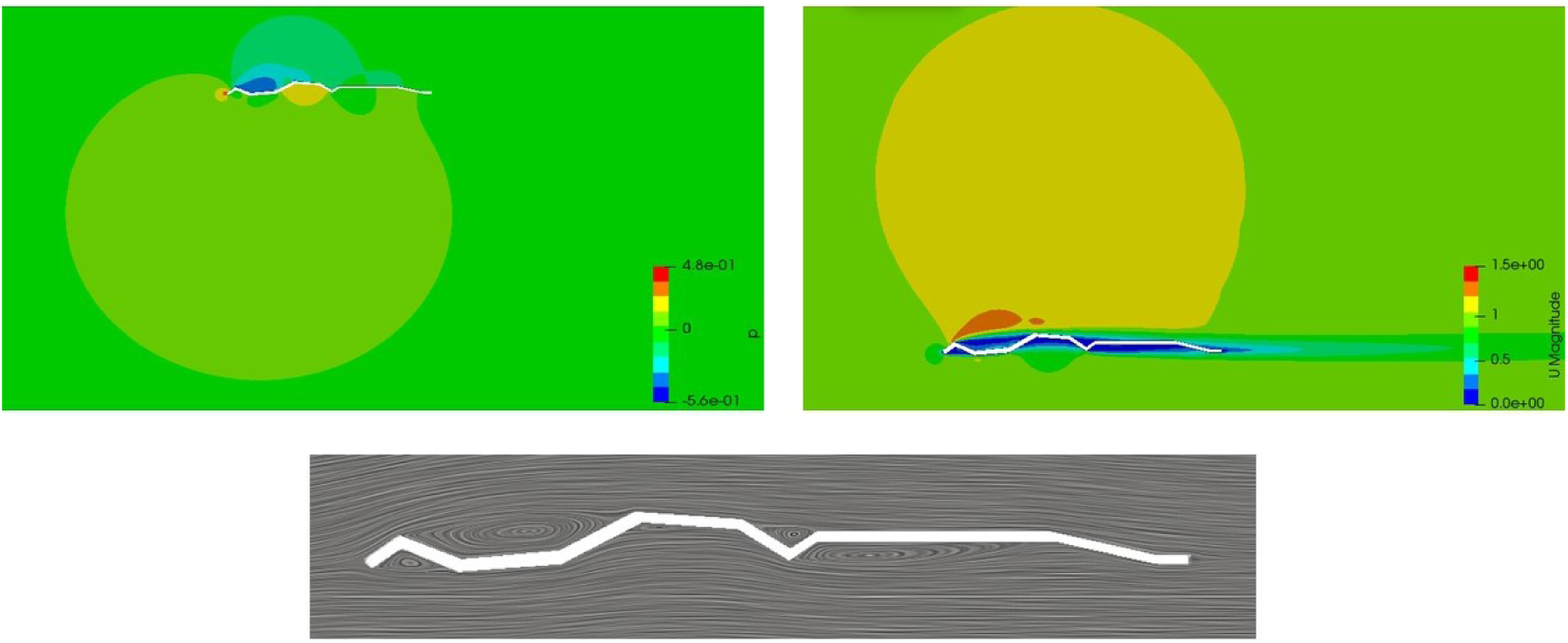
Up, left: pressure field. Up, right: velocity field. Bottom: streamlines of the vortices. Results referred to profile 3, in correspondence of an AoA = 2°

### 3.2. Results

The obtained results in terms of lift and drag coefficients are compared with those obtained by Kesel [11]. The graphs in figure 18 clearly display the similarity between the results of these numerical analyses and the outcomes of the experimental tests made by [11] on profile 2.

**Figure 18:**
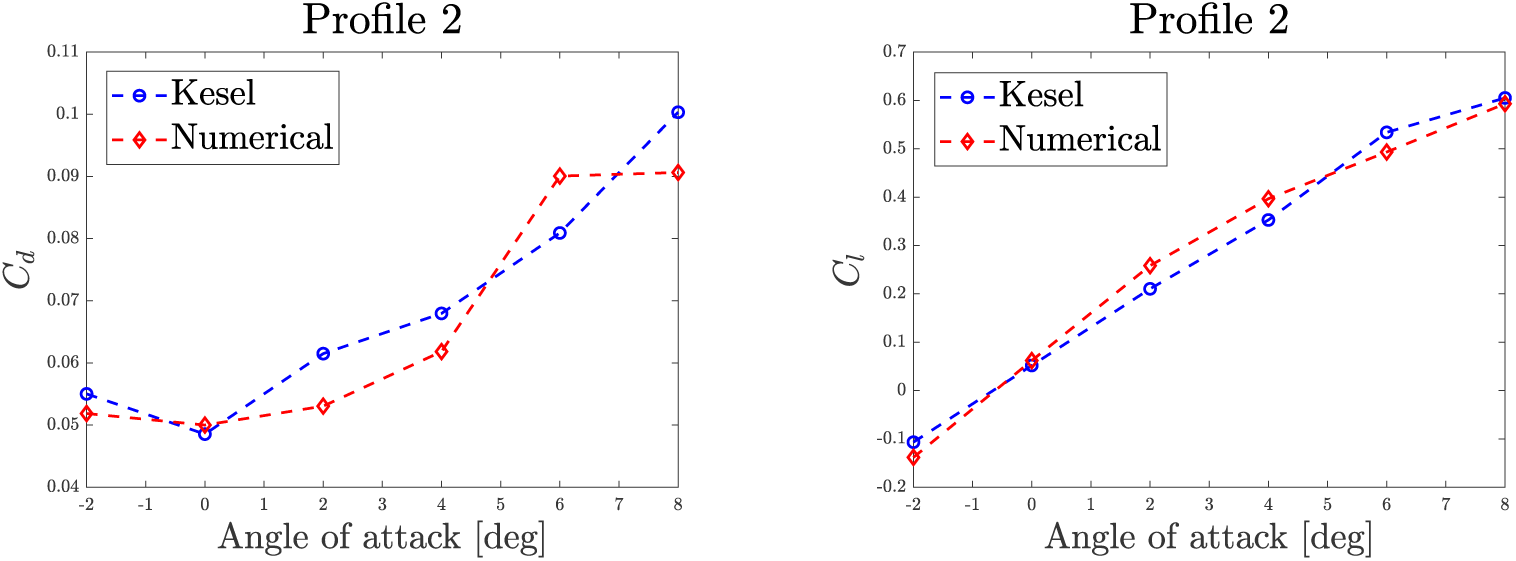
Comparison of lift and drag coefficients obtained with these simulations with the experimental results of Kesel [11]

From simulations of profile 2 a very high level of correspondence between numerical and experimental results emerged. This can be explained by the fact that the less the inclination of the leading edge, the lower the magnitude of the vortex-detachment phenomenon in correspondence of the front part of the wing, where the first contact between the airflow and the body takes place. Less flow separation means that unsteady effects, which are disregarded by a steady-state solver, are less important, and so a better characterisation of the phenomenon could be achieved. For the other two profiles this does not hold as well, therefore such similarity cannot be expected, also because near wing root and wing tip the effects of a 3D flow are less negligible. Nevertheless, the same trend in both drag and lift coefficient is pointed out for all the angles of attack, as shown in figures 19 and 20.

**Figure 19:**
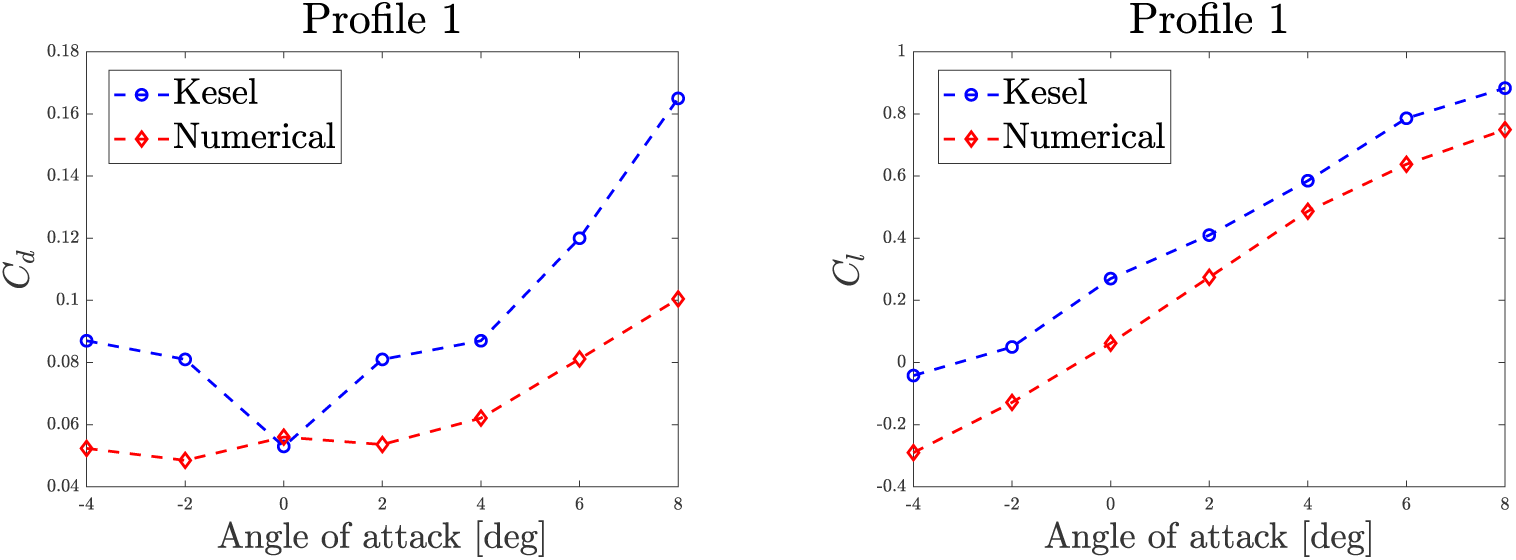
Comparison of lift and drag coefficients obtained with these simulations with the experimental results of Kesel [11] on profile 1

**Figure 20:**
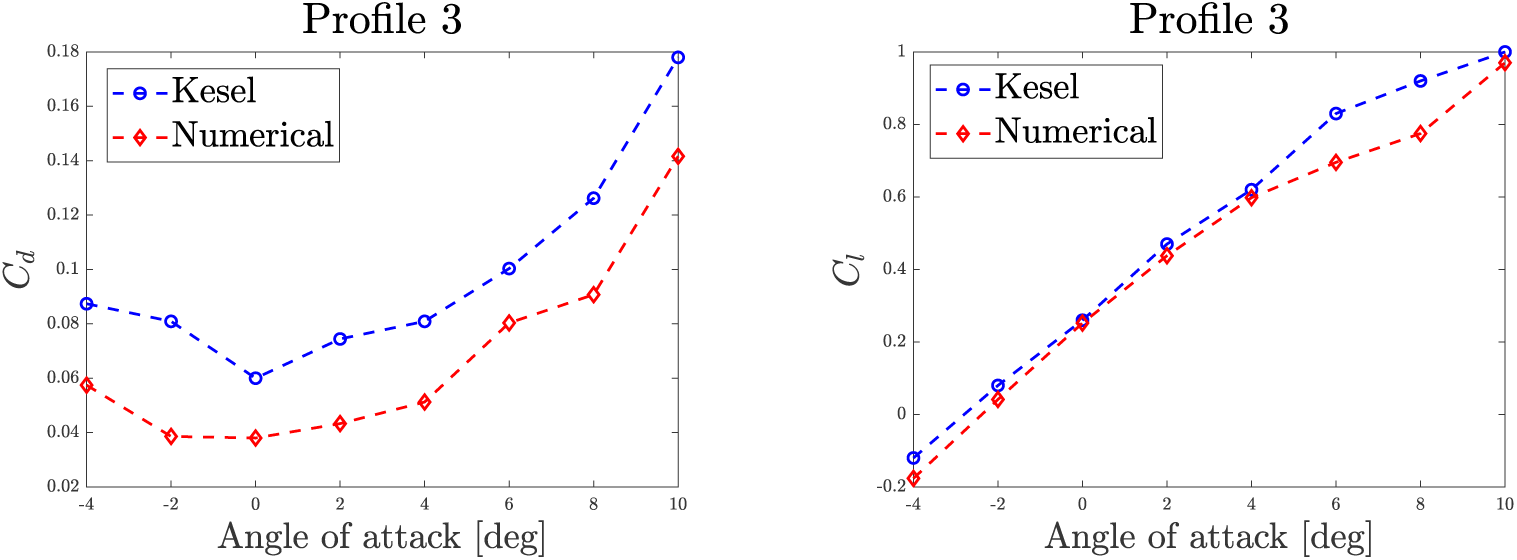
Comparison of lift and drag coefficients obtained with these simulations with the experimental results of Kesel [11] on profile 3

As it can be noticed, looking at figures 18, 19 and 20, the accuracy of the model proved to be slightly different among the profiles. However, the capability of the numerical model to represent the trends pointed out by the experiments is assessed, although many simplifying hypotheses have been made to reduce the degree of complexity of the model, focusing only on the core characteristics of the phenomenon.

## 4. Hovering: numerical analysis

Having determined the aerodynamic coefficients of three sections of a dragonfly *Aeshna Cyanea* forewing, it has been deemed interesting to verify the capability of the flyer to keep itself aloft. This kind of flight condition is called hovering: the insects flaps its wings, despite remaining still in the air. Translating such maneuver into dynamics, the objective of the numerical analysis could be expressed as a verification that the wings engender enough lift to sustain the insect’s weight. Hovering is correctly represented if the horizontal force - thrust -is null, while the vertical one -lift -equals the weight of the insect. For this analysis the 4 wings are considered as equal, thus calculating the total lift multiplying times 4 the vertical force generated by the single wing. Nevertheless, such hypothesis should lead to conservative results, indeed Xie and Huang (2015) [7] showed that the pair of hindwings is able to engender higher lift force than the forewings.

Recognised the similarity of the topic with the realm of wind turbines, a methodology inspired to the Blade Element Momentum (BEM) theory is applied. Such procedure involves breaking a blade down into several small parts then determining the forces on each of these small blade elements. These forces are then integrated along the entire blade over one period in order to obtain the forces and moments produced by the entire body. Analogously, the dragonfly wing has been divided into 10 regions, 5 mm each, approximated to rectangles whose length is equal to the mean chord of each section. All the geometrical dimensions needed to perform the simulations have been taken from the article by Kesel [44]. The contribution of the body is usually very small for insects. As a matter of fact, the study by Yao and Yeo (2017) [3] about the flight of the *Drosophila Melanogaster* proved that the influence of the body on the generation of the forces never overcomes the 5%, no matter the flight regime. More specifically, in normal hovering such contribution is of the order of 1%, thus, if neglected, not invalidating the goodness of the simulation. The following hypoteses about *C*_*L*_ and *C*_*D*_ for each region were made:

- **sections 1 to 3**: *C*_*L*_ and *C*_*D*_ coefficients assigned as the ones calculated for profile 1;
- **section 4**: *C*_*L*_ and *C*_*D*_ coefficients assigned as the mean value of the ones calculated for profile 1 and 2;
- **sections 5 to 6**: *C*_*L*_ and *C*_*D*_ coefficients assigned as the ones calculated for profile 2;
- **section 7**: *C*_*L*_ and *C*_*D*_ coefficients assigned as the mean value of the ones calculated for profile and 3;
- **sections 8 to 10**: *C*_*L*_ and *C*_*D*_ coefficients assigned as the ones calculated for profile 3.

In accordance with the majority of articles about this topic ([[27]], [[19]]) for each section and for each time instant, velocity and angle of attack are computed considering the following motion law (figure 22):

**Figure 21:**
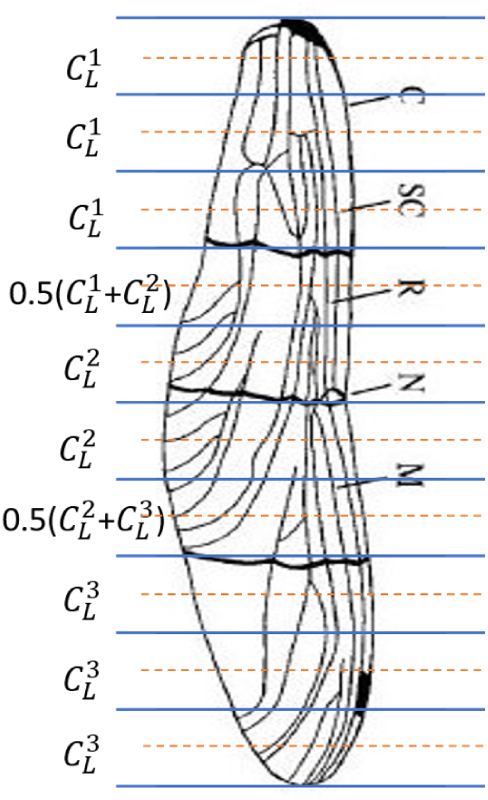
Subdivision in sections of the dragonfly’s wing and correspondent lift coefficients

**Figure 22:**
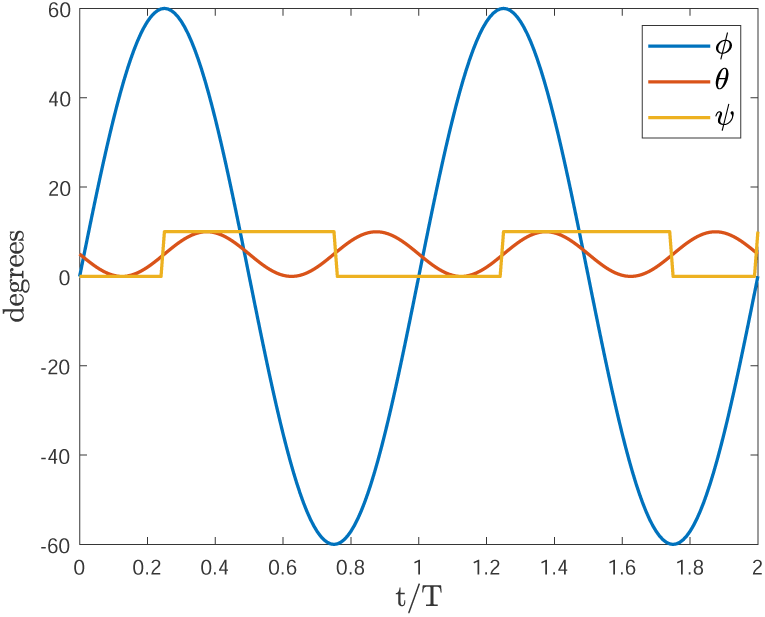
Motion laws imposed to the angles of the wing during hovering simulation

- Sweep angle, *ϕ*: sinusoidal. Amplitude: 60°, frequency: 35 Hz, phase: 0°;
- Elevation angle, *θ*: 5° upward-shifted sinusoidal. Amplitude: 5°, frequency: 70 Hz, phase: 180°;
- Twist angle, *ψ*: 5° upward-shifted square wave. Amplitude: 5°, frequency: 70 Hz, phase: 0°.

The outcomes are presented under the form of a mass value, in milligrams, that the flyer proved to be able to displace along two directions: horizontal, and vertical. As far as the horizontal carrying capability is concerned, the outcome is of 17 mg, whereas 998 mg is the vertical carrying capability as resulted from this simulation. According to the research by Grabow and Ruppell (1995) [29], the average weight of an *Aeshna Cyanea* depends on the gender: 729 mg for the males, 1098 mg for the females. Both the values fit very well with the results of the simulation. Regarding the horizontal payload capability, the result represents less than 2% of the weight of the insect, thus deemed to be a satisfying outcome.

## 5. Conclusions

An overview of the most important kinematic and biological parameters affecting the dynamics of insects’ flight is presented in the initial part of this article. The focus has been set on the dragonfly, whose flight characteristics were deemed promising for as regards the design of MAVs. From then on the article is hinged on numerical studies. First, the results of the comparison between a simplified CFD model and an experimental study [11] focused on 2D gliding flight are shown. The numerical model proved to be accurate and able to describe the micro-turbulent nature of the flow in which the dragonfly is immersed. The impressively good values of aerodynamic coefficients attained by the profiles object of simulation confirmed the suitability of the dragonfly as a source of inspiration for the design of MAVs. Conclusively, a 3D hovering simulation has been conducted in order to study a peculiarity of the kinematics of such flyer: the “8-figure” wingbeat. To do so, instead of exploiting a three-dimensional CFD code, it has been decided to apply the BEM methodology, well established in the realm of the design of wind turbines. The model behaved coherently with the flight condition that was represented, although the novelty of such application and the approximations that were made. Looking at the work as a whole, it presents to the reader the design of two numerical tools for the analysis of the forces generated by a generic body involved in a low-Reynolds flight regime, enriched by a synthesis of the effect of some relevant kinematic and biological parameters on the dynamics. Despite in this article the numerical analysis has been applied to the specific case of the dragonfly, it is possible to adapt the models to study different wings in a wide variety of flight conditions, realized by changing the parameters listed in section 2. In other words, one may decide to model the body to examine and its kinematics on the basis of the variables presented in the literature review, and then inspect their effects on the engendered forces exploiting the numerical analyses presented in the following two sections. Conclusively, such study could widen the horizons of the design of MAVs providing the possibility of testing a large variety of configurations.

